# FLONE: fully Lorentz network embedding for inferring novel drug targets

**DOI:** 10.1101/2023.03.20.533432

**Authors:** Yang Yue, David McDonald, Luoying Hao, Huangshu Lei, Mark S. Butler, Shan He

## Abstract

**Motivation:** To predict novel drug targets, graph-based machine learning methods have been widely used to capture the relationships between drug, target, and disease entities in heterogeneous biological networks. However, most of the existing methods cannot explicitly consider disease types when predicting targets. More importantly, drug-disease-target (DDT) networks could exhibit multi-relational hierarchical sub-structures with the information of biological interactive functions, but these methods, especially those based on Euclidean space embedding cannot fully utilize such topology information, which might lead to sub-optimal prediction results. We hypothesized that, by properly importing hyperbolic space embedding specifically for modeling hierarchical DDT networks, graph-based machine learning algorithms could better capture the relationships between aforementioned entities, which ultimately improves the target prediction performance.

**Results:** To test our hypothesis, we formulated the drug target prediction problem as a knowledge graph triple completion task explicitly considering disease types. To solve this problem, we proposed FLONE, a novel hyperbolic Lorentz space embedding-based method to capture the hierarchical structural information in the DDT network when inferring drug targets. We evaluated it on two different DDT networks, our experimental results showed that by introducing Lorentz space, one of the isomorphic models of hyperbolic space, FLONE generates more accurate candidate target predictions given the drug and disease than the Euclidean translation-based counterparts, which supports our hypothesis. Beyond these experiments, we also devised the hyperbolic encoders to fuse drug and target similarity information into FLONE, to make it capable to handle triples corresponding to previously unseen drugs and targets for more challenging prediction scenarios.

**Availability and implementation:** Source code and dataset information are at: https://github.com/arantir123/DDT_triple_prediction.

## 1 Introduction

Inferring novel drug targets based on computational methods has attracted more attention recently, because it can effectively reduce the time cost in the early stages of drug development (Oprea, et al., 2011). An important systematic strategy for inferring drug targets is based on analyzing known relationships between drug, target, and disease entities in biomedical databases, e.g., DrugBank (Wishart, et al., 2006) and Pharos (Nguyen, et al., 2017). The known relationships in these databases are usually constructed as knowledge graphs (KGs) or heterogeneous biological networks and then are analyzed by graph learning algorithms (Bordes, et al., 2013; Grover and Leskovec, 2016). For example, Luo, et al. compiled their dataset DTINet which contains drug-disease-target (DDT) relationships from DrugBank, CTD (Davis, et al., 2019), HPRD (Keshava Prasad, et al., 2009), and SIDER (Kuhn, et al., 2010) databases, and the network diffusion algorithm and inductive matrix completion strategy were used to infer novel drug-target interactions (DTIs) (Luo, et al., 2017). Peng, et al. developed a heterogeneous graph convolutional neural (GCN) network to learn multi-relational information in DDT-relevant networks for DTI predictions (Peng, et al., 2021).

However, these methods usually predict putative associations between drugs and targets, ignoring inferring the causal relationships between diseases and drugs/targets. For example, in DTINet, only 58 out of 5603 diseases have modulation relationships with the target *RBP3* (*P10745*) of drug *Vitamin A* (*DB00162*), in this case, explicitly considering disease types could bring finer scale virtual screening, allowing models to predict drug targets within particular disease types.

Furthermore, most existing methods modelled target predictions as a (relatively balanced) binary classification task, i.e., whether there is an interaction between a drug-target pair. For training and evaluating the binary classification model, apart from the positive samples (known DTIs), both training and test sets are supplied with a fixed number of negative samples by randomly selecting from all candidate targets (e.g., 1:1 with positive samples). In this case, the performance evaluation reflects how well the model distinguishes positive samples from a manually selected subset of candidate targets that could have bias.

To address these problems, Moon, et al. formulated the target prediction as a recommendation system ranking task (Moon, et al., 2021). The ranking task aims to directly assign each candidate target in the network an interaction probability score for the given drug and disease. The scores should allow positive targets to rank higher than negative candidate targets. This ranking task formulation better evaluates the model’s capability to identify positive target samples from all candidate targets, which better reflects true model performance in actual DTI virtual screening.

However, the method DDTE in Moon, et al could be further improved by capturing the intrinsic hierarchical structure (i.e., the tree-like structure) in the DDT networks. As shown in Figure 1, the complex interactions in a DDT network are hierarchically organized. For example, for a drug node *DB00050* in DTINet, which can be seen as a root node, we know it binds to two target nodes *P30968* and *P22888* directly, with which associate 110 different types of disease nodes based on drug-disease and disease-target edges. These drug-target-disease relationships essentially form a hierarchical structure, which provides extra topological information about the interaction mechanism of the DDT network. Capturing this hierarchical structure information in the DDT network could be helpful to generate more accurate predictions.

**Fig. 1.**
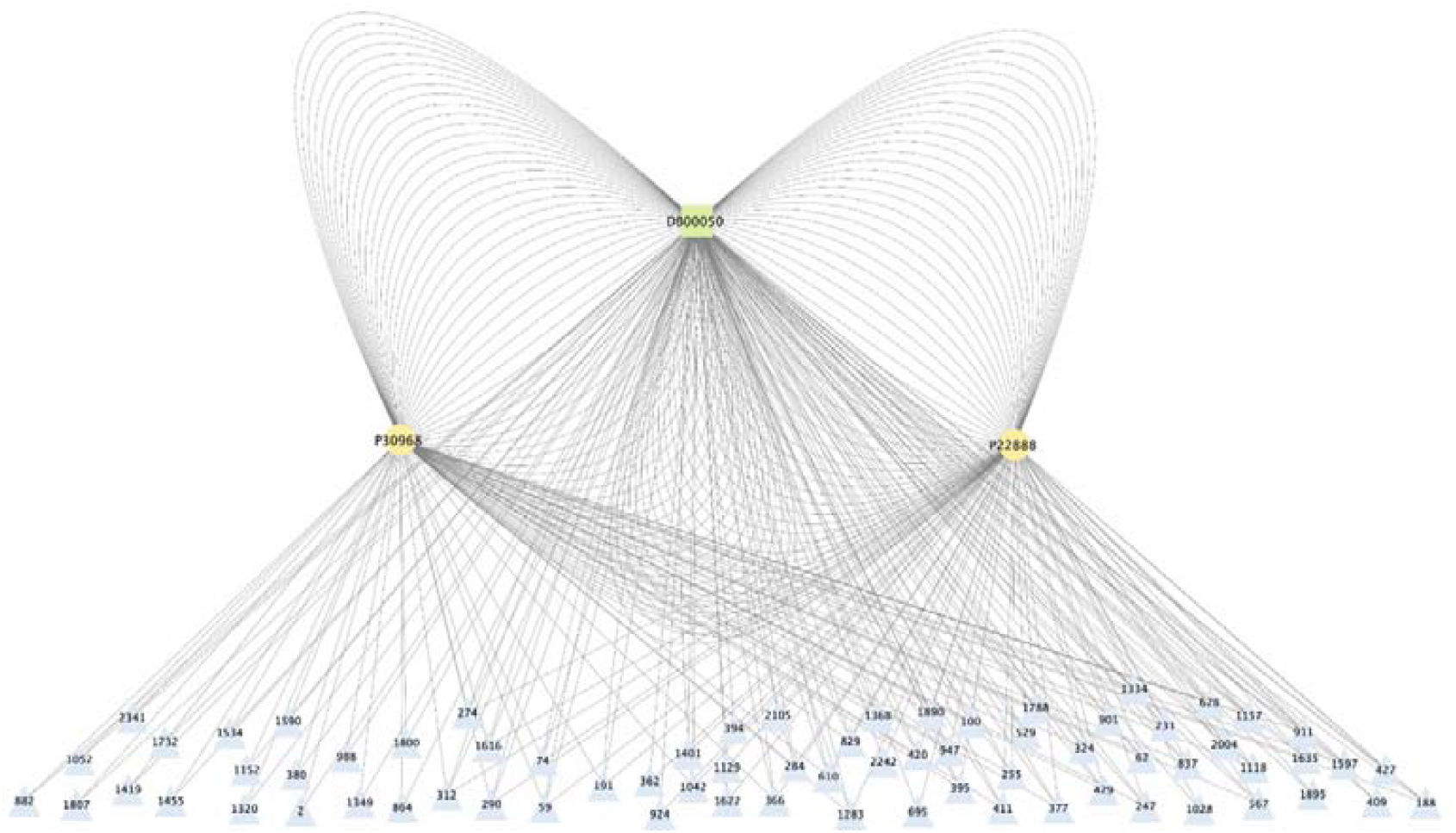
Illustration of the hierarchies in the DDT network. Taking drug node *DB00050* in DTINet as an example, it can bind to two target nodes *P30968* and *P22888* directly, it can also associate with these two targets through 110 different types of disease nodes based on drug-disease and disease-target edges (the labels of disease nodes are disease ids used in FLONE). Thus, starting from *DB00050*, a local hierarchy can be formed, and such structure can be generalized to other drugs, forming more complicated implicit hierarchies that reveal the interactive mechanism of DDT networks.

Nevertheless, despite the importance, most of the graph learning methods including DDTE cannot effectively capture this hierarchical information (of the DDT network) because they work in Euclidean space. It is known that Euclidean space grows polynomially, limiting its capacity to represent hierarchical networks in which the volume of the hierarchies increases exponentially in proportion to its radius (Yang, et al., 2022). In contrast, hyperbolic space, which can be seen as a continuous tree space, is a better alternative than Euclidean space since it can fit hierarchies with its exponentially increasing capacity and smaller distortion (Chen, et al., 2021). Based on this, we hypothesized that, by properly introducing hyperbolic space, graph machine learning algorithms could better capture the implicit hierarchies in the DDT networks, generating more accurate predictions for every candidate target (under explicitly given drugs and diseases). To test this hypothesis, we first formulated the drug target prediction as a (hyperbolic) translation-based knowledge graph drug-disease-target triple completion (KGC) problem explicitly considering disease types, in which drugs and diseases are subjects and predicates separately, and targets are the objects to be completed (i.e., predicted).

To solve this problem, we proposed FLONE (fully Lorentz network embedding), in which the main component is a hyperbolic similarity calculation module based on the fully Lorentz linear transformation (FuLLiT) (Chen, et al., 2021). FuLLiT essentially calculates the Lorentzian distance (i.e., similarity) between the hyperbolic embeddings of candidate targets and the hyperbolic representation of the given drug and disease. The Lorentzian distance is then used to infer each candidate target of a given drug and disease. Unlike other Euclidean embedding methods, FuLLiT can capture the implicit hierarchies by learning our defined drug-disease-target triple sets extracted from the DDT network.

Another important feature of FLONE is its ability to handle previously unseen drug and target entities when embedding drug-disease-target triples, which is critical when applied to real-world problems. Our idea to handle unseen drugs and targets is to use the hyperbolic drug and target encoders based on the fully Lorentz linear and fully Lorentz GCN layers, to fuse drug and target similarity information into the learning process of the hierarchical structure.

Within the scope of the aforementioned practical application scenario (i.e., explicitly considering disease types and directly ranking all candidate targets), we conducted extensive experiments on FLONE to test our hypothesis. Our study showed that the translation-based drug-disease-target embedding benefits from the Lorentz space, which supports our hypothesis. Apart from demonstrating this hypothesis, our results also showed that by fusing the drug structure and target sequence similarity as additional domain knowledge, FLONE not only achieved better predictions on drug-disease-target triples related to seen drugs and targets, but also could provide accurate predictions on those corresponding to previously unseen drugs and targets.

## 2 Materials

### 2.1 Datasets

To construct the heterogeneous drug-disease-target networks based on the DTINet (Luo, et al., 2017) and BioKG (Walsh, et al., 2020) datasets, we first defined the extraction rule of the drug-disease-target triple set. Specifically, we extracted a triple (*drug*_*i*_ *diseasek*_*k*_, *target*_*j*_) (abbreviated as 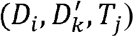) if there is a positive/known edge between every two of three of *drugt*_*i*_, *target*_*j*_, and *disease*_*k*_ in DTINet and BioKG (i.e., all the three edges are known). This stringent extraction rule ensures that all three of the 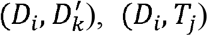, and 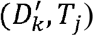 pairs exist, which improves the quality of the extracted triples.

However, in KGC tasks, there could be implicit data leakage caused by very similar predicates, e.g., for two *(subject, predicate, object)* triples, *(‘Birmingham’, ‘is_in’, ‘UK’)* and *(‘Birmingham’, ‘is_located_in’, ‘UK’)*, because the predicates ‘*is_in’* and *‘is_located_in’* have very similar semantic meaning, thus they correspond to very similar subject-object pairs, which causes over-idealistic results when predicting triples related to ‘*is_in’* or *‘is_located_in’*. To avoid this problem, we removed diseases that had >60% drug-target pair similarity with other diseases based on the Jaccard similarity coefficient, which generated 171597 positive tuples consisting of 535 drugs, 417 targets, and 1160 diseases from DTINet as well as 9699 positive triples consisting of 1128 drugs, 723 targets, and 529 diseases from BioKG. Furthermore, unlike the common triple setting: for a 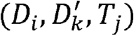 tuple, *D*_*i*_, *T*_*j*_, and 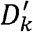 represent the drug node, target node, and disease edge between them, respectively; in our task, these three items are all nodes in DDT networks, thus what FLONE predicts is a novel triangular (relationship) sub-structure between them. However, inappropriate data splitting for model evaluation will lead to another type of data leakage, which is detailed in the Section 4.2.

## 3 Methods

### 3.1 The basic definition of the Lorentz model

The hyperbolic space is defined as a smooth Riemannian manifold equipped with the constant negative curvature and positive-definite inner product on the tangent space at every point. There are several isomorphic geometric models of the hyperbolic space:

Lorentz model, Poincaré disk model, Poincaré half-plane model, and Klein model. In this paper, we used the Lorentz model, which is one of the widely used models because of its numerical stability during training and closed-form computation of geodesics (Chen, et al., 2021; McDonald, et al., 2022; Sun, et al., 2021; Wu, et al., 2021; Yu, et al., 2020).

A *n*-dimension Lorentz model is defined as 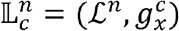, where *c* is the constant negative curvature, 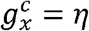 is the Riemannian metric satisfying *η* = *I* (*I* is the (*n* + 1) -dimension identity matrix) except *η η*_0,0_ = −1, and ℒ^*n*^ represents the coordinates/features of the point set in 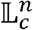:

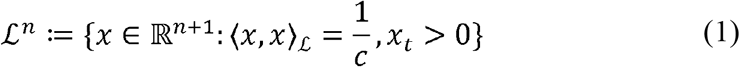

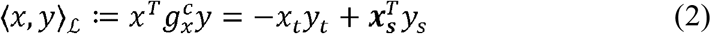

where ⟨…,…⟩_ℒ_ is the Lorentzian inner product. Each point *x* in 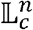 (i.e., 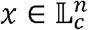) is expressed as a concatenation [*x*_*t*_, *x*_*s*_], where *x*_t_ ∈ ℝ (referred as the time dimension), *x*_*t*_ ∈ ℝ^*n*^ (referred as the spatial dimension), and the origin 𝒪 of the Lorentz model is defined as 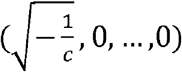.

Besides, for every point 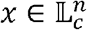, it is equipped with an orthogonal space (i.e., tangent space) of 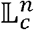 (at *x*), which is the first-order approximation of ℒ^*n*^ around *x* (Yang, et al., 2022), and is formally defined as 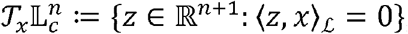, where, is the point set of this tangent space.

### 3.2 Description of the FLONE method

Based on the defined Lorentz model, we proposed FLONE to solve the drug-disease-target triple target entity completion problem. Specifically, for each drug-disease-target triple in the extracted triple set, FLONE treats the drug, disease, and target as the subject, predicate/relation, and object, respectively. The task is, given a drug-disease combination, FLONE will assign a similarity score to every candidate target (i.e., object entity) in the DDT network, which indicates the distance similarity between the target and the given drug and disease. FLONE then uses the similarity scores to rank these targets, for selecting high-confidence targets for the current drug-disease combination.

In other words, a high-quality model would directly assign high similarity scores to all of the defined drug-disease-target triples that have been extracted from the DDT network, and small similarities for all other drug-disease-target combinations. A researcher investigating drug repurposing would provide the drug and disease of interest and will have a ranked list of targets returned for further investigation.

The illustration of FLONE is shown in Figure 2. After the extraction of triples from the heterogeneous DDT network, we presented FLONE as an end-to-end framework consisting of three major components:

**Fig. 2.**
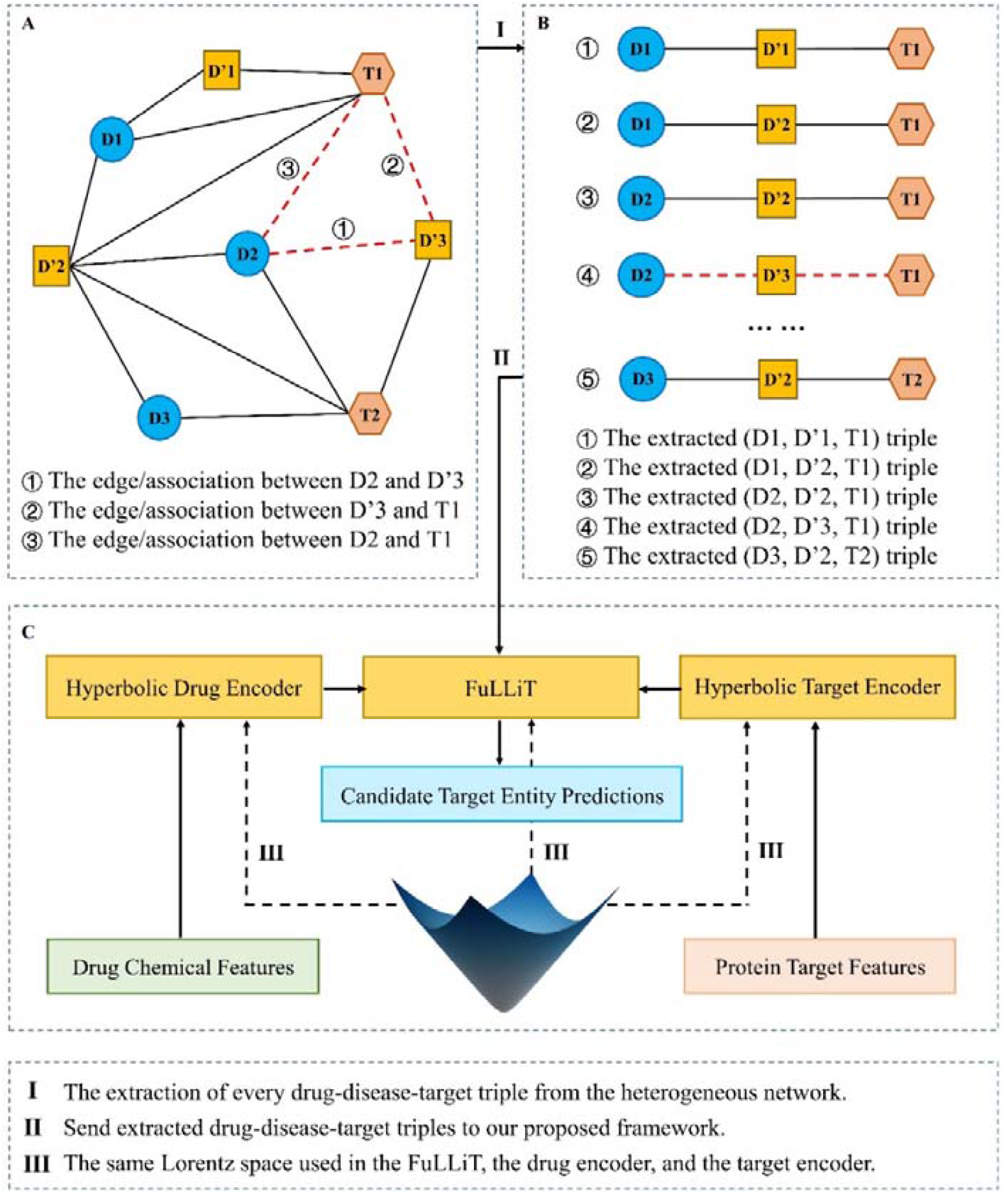
Illustration of FLONE. (A) Construct a heterogeneous DDT network. (B) Decompose this network to extract the drug-disease-target triple set. (C) Input the Euclidean drug and target domain knowledge into the hyperbolic drug and target encoders respectively, to generate hyperbolic drug and target embeddings (or using self-contained drug and target embeddings in FuLLiT directly). Next, the hyperbolic drug and target embeddings are used by FuLLiT, in which hyperbolic drug embeddings are transformed by the disease translation representations, and then the Lorentzian distance-based similarity between the transformed drug embeddings and hyperbolic target embeddings are calculated. After training, FLONE is able to rank every candidate target entity for the given drug and disease.

1. FuLLiT, which was proposed to increase the performance of the Euclidean translation-based KGC method in our task. FuLLiT treats a predicate/relation as a trainable translation/representation offset from the subject to the object entity embeddings. It takes the drug and target hyperbolic embeddings and a disease as inputs, and outputs a similarity score for the current drug-disease-target triple. Besides, it works on the Lorentz space with the fixed dimension and constant negative curvature (−1 in our study). We discussed it in the Section 3.3.
2. Hyperbolic drug encoder, which is responsible for taking drug chemical features to produce a hyperbolic embedding for every drug in the extracted triple set. It can inject drug domain knowledge into FuLLiT for handling previously unseen drugs (explained in more detail in the Section 3.4).
3. Hyperbolic target encoder, which, likewise, produces hyperbolic embeddings for all of the targets extracted from the DDT network, for injecting target domain knowledge into FuLLiT for handling unseen targets (detailed in the Section 3.4).

### 3.3 Description of FuLLiT

All important (hyperbolic) computation operations of FLONE are fully Lorentzian instead of a hybrid mode (Chen, et al., 2021). This has the advantage that no need to expensively map between the Lorentz space and its tangent space (McDonald, et al., 2022). Based on the fully Lorentz linear layer, the fundamental module of FLONE – FuLLiT was developed, which includes three components: a hyperbolic triple decoder, a self-contained drug embedding look-up table, and a self-contained target embedding look-up table. The first component is to calculate the similarity score for the drug-disease-target triples, and the latter two are for providing corresponding hyperbolic embeddings without domain knowledge. These three components form the backbone of FLONE that can deal with cases where external domain knowledge is unavailable.

Specifically, the general form of the fully Lorentz linear layer *FLLinear*_*n,m*_ *(x)* is defined as (3), which could ensure a linear transformation to map 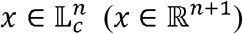 to 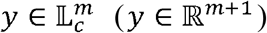 (i.e., the coordinates of input x and output y of the transformation are guaranteed to stay in the respective Lorentz space):

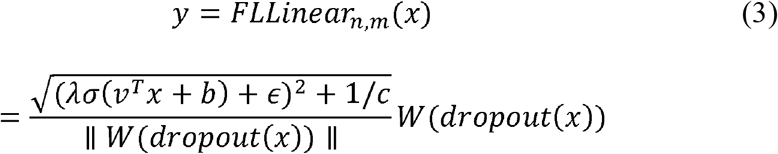

where *λ* is a fixed hyper-parameter to control the numerical scale of the time dimension, *σ* is the sigmoid function, *v* ∈ ℝ^*n*+1^ and *W* ∈ ℝ^(*n*+1)^ are trainable weights of the transformation matrix 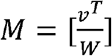 in this linear layer. Besides, b is the trainable bias, is a fixed value larger than 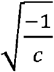 to ensure 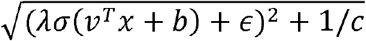 larger than 0. In FLONE, we adopted the same dimension for all embeddings, in this case, *n* is equal to *m* in every intermediate *FLLinearn*_*n,m*_ (*x*) layer.

For the hyperbolic triple decoder, assuming embedding drug and target entities into a *n*-dimension Lorentz space, when calculating similarity probability scores (from 0 to 1), for example, to predict the score for *T*_*j*_ under 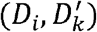, five sets of parameters including corresponding drug and target embeddings, disease type specific translation offset, and real-value drug and target biases are needed. Specifically, we first obtained the hyperbolic embedding 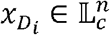 and 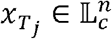 of *D*_*i*_ and *T*_*j*_ from the self-contained embedding look-up tables (i.e., the tables that store the corresponding type of embeddings through entity indices). Then, 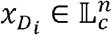 was transformed by the disease type specific (i.e., 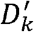 in this case) *FLLinear*_*n,n*_(*x*), to obtain the translation 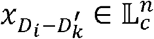. Then the similarity probability score p can be calculated as (Balazevic, et al., 2019; Chen, et al., 2021):

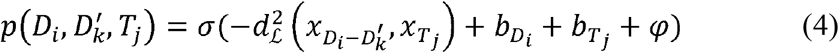

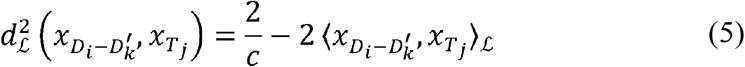

where 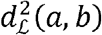 is the squared Lorentzian distance, 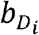 and 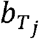 are the drug and target type specific biases, respectively. *φ* is the margin hyper-parameter, and *σ* is the sigmoid function.

Furthermore, the self-contained drug and target embedding look-up tables are essentially trainable matrices with the (*entity number, embedding size*) shape, trained together with other sets of parameters, to produce the required hyperbolic embeddings based on learning topological structure information of DDT networks. Besides, the look-up tables can also be created by hyperbolic drug and target encoders (detailed in the next section) in an end-to-end way (i.e., all sets of parameters are learnt simultaneously based on known triples extracted from the DDT network).

As a comparison, the representative Euclidean translation-based method DDTE (Moon, et al., 2021) mentioned in Introduction uses vectors as disease translation offsets, i.e., the translated/transformed drug embedding is calculated by 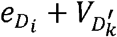 where 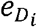 is the Euclidean drug embedding and 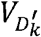 is the vector-based disease offset (while the matrix-based offset 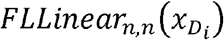 is used in our FuLLiT). Except for specialized hyperbolic operations, another two main differences between DDTE and FuLLiT/FLONE are that, compared with FLONE, when calculating the similarity probability score, there are no drug and target specific biases of formula (4) in DDTE, and DDTE cannot consider any external domain knowledge (i.e., only providing the Euclidean self-contained drug and target look-up tables).

### 3.4 The hyperbolic encoders for incorporating drug and target similarity information

To investigate whether the triple target entity completion benefits from the fusion of domain knowledge, we extended the FuLLiT, by replacing the self-contained drug and target embedding look-up tables with the tables generated by corresponding hyperbolic encoders, for encoding and injecting external domain knowledge into embeddings, and the size of these embeddings is same as that in the self-contained look-up tables.

Specifically, we tried to fuse different domain knowledge of drug-drug and target-target similarities by using the hyperbolic drug and target encoders. Intuitively, this could bring similar hyperbolic embeddings for similar drugs/targets, which further improves the model’s predictive performance (Luo, et al., 2017). For the drugs in the network, we collected ECFP6, a powerful circular topological fingerprint commonly used in drug discovery (Rogers, et al., 2010), as the input of the hyperbolic drug encoder. For involved targets, we provided two efficient target features to be encoded by the hyperbolic target encoder: graph-like features (i.e., GO-PPI network) and sequence-like features (i.e., target sequence similarity). A detailed description of constructing the GO-PPI network is in the Supplementary Material.

To encode these (Euclidean) drug and target prior features, the tangent space is needed to map them into the Lorentz space. Specifically, for the hyperbolic drug encoder, it takes the (Euclidean) ECFP6 as the input. Take encoding ECFP6 of *D*_*j*_ (terms as 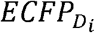) as an example, first, 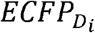 is concatenated with 0 to create 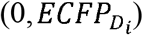, to map 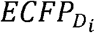 into the tangent space of 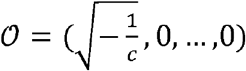. Because according to 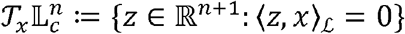 and 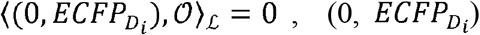 is in the tangent space of 𝒪. Then, 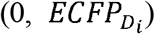 is further mapped into the Lorentz space through the exponential map function as follows, where 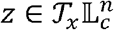:

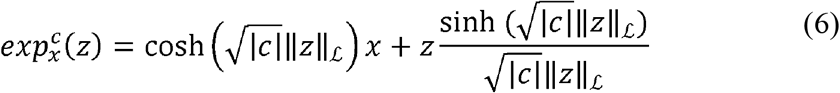

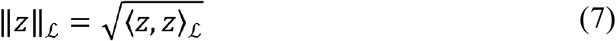

In this way, the Lorentzian ECFP6, i.e., 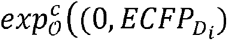, can be generated, and then 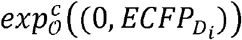, is sent to *FLLinear*_*m,n*_(*x*) specifically for encoding Lorentzian ECFP6, to reduce its dimension to the unified hidden dimension mentioned in the Section 3.3. Furthermore, all Lorentzian ECFP6 (after dimension reduction) of involved drugs constitutes the hyperbolic drug embedding look-up table (through drug entity indices). As an analogy, using the hyperbolic target encoder for the sequence-like target features (i.e., target sequence similarity in our experiments), the hyperbolic target embedding look-up table can be generated. In addition, the description of generating the hyperbolic target embedding look-up table based on GO-PPI network is provided in the Supplementary Material.

After introducing the hyperbolic encoders to FuLLiT, the parameters to be end-to-end optimized are not the weight of self-contained embedding trainable matrices anymore but different weights in fully Lorentz layers (of the encoders). The advantage of adding domain knowledge-based encoders is that, previously unseen drugs and targets can be handled by the injected prior similarity, and the encoder architecture can be made arbitrarily deep/complex as the researcher desires.

### 3.5 Model training and optimization

Similar to the Euclidean translation-based KGC models, negative sampling (Bordes, et al., 2013) was used to train the FLONE. Specifically, during the training phase, for each positive/known triple 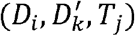, a set of negative triples were generated by randomly replacing *T*_*j*_ with other *T*_*j*′_ in the target entity set and ensuring that the generated negative triples were not in the extracted drug-disease-target triple set.

The loss function used for optimizing all sets of parameters in FLONE was the binary cross entropy (BCE) loss defined as follows:

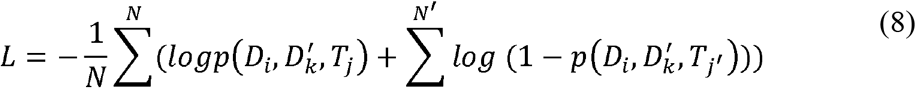

where *N* is the positive/known triple number (in the training set) and *N*′ is the number of negative triples generated for each known triple. Besides, we adopted the Riemannian Adam as the optimizer of FLONE, which is the counterpart of Adam defined in Euclidean space (Bécigneul and Ganea, 2018; Kochurov, et al., 2020).

## 4 Results and discussion

### 4.1 Hyperbolicity test of the DDT networks

To test whether the resulting heterogeneous DDT networks (equivalent to the extracted triple sets) exhibit a hierarchical structure for demonstrating our hypothesis, we calculated their Gromovs hyperbolicity *δ* (Chami, et al., 2019; Gromov, 1987), which measures how hierarchical the network is. The lower *δ*, the more implicit hierarchies the network has, and *δ* of completely tree-like structures is 0. Moreover, for common hierarchical benchmark datasets, e.g., Human PPI and Airport, their *δ* is about 1, and for the standard (non-hierarchical) benchmark, e.g., Cora, its *δ* is 11 (Chami, et al., 2019). While the calculated *δ* values of the extracted DTINet and BioKG DDT networks were both 1.5, which indicated that these networks do possess implicit hierarchies, making them theoretically feasible for hyperbolic space embedding.

### 4.2 Model evaluation settings

Except for the data leakage mentioned in the Section 2.1, the common model evaluation setting, i.e., randomly splitting the drug-disease-target triple set into training, validation, and test sets could also cause over-idealistic results in our task. This is because each defined positive/known 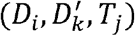 triple not only contains the associations 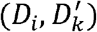 and 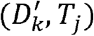, but also includes (*D*_*i*_, *T*_*j*_) (i.e., constituting a triangular inter-node sub-structure), and ignoring (*D*_*i*_, *T*_*j*_) in data splitting will lead to extra data leakage. For example, if 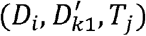 and 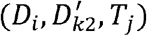 are allocated into training and test sets separately, after training, when inferring the target for 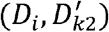, *T*_*j*_ will be chosen more easily than other candidate targets, as the model ‘has already seen’ the unnecessary implicit association information between *D*_*i*_ and *T*_*j*_ (i.e., (*D*_*i*_, *T*_*j*_)) during the training phase.

To avoid this pitfall, we split these triples based on drug-target pairs. In other words, the known triples with the same drug-target pair were put into the same set (i.e., training, validation, or test sets). In this case, a part of the drugs and targets in the test set did not occur in the training set (the related statistics are in the Supplementary Material), which also increased the difficulty of target predictions. Furthermore, the targets to be inferred in the test set must be previously unseen for the given drug.

Based on this setting, the drug-disease-target triples corresponding to 60%: 20%: 20% of all drug-target pair varieties were divided into training, validation, and test sets, respectively. This procedure was repeated five times independently, for each time, before splitting data, the whole drug-disease-target triple set was randomly shuffled to make different drug-target pair varieties enter each set. We computed and reported the average evaluation metrics over the five independent repeats.

To evaluate the model’s predictive performance in the scenario of explicitly considering disease types and directly ranking all candidate targets, we adopted the standard evaluation metrics used in recommendation system ranking tasks, including Mean Reciprocal Rank (MRR), Hits@1, Hits@3, and Hits@10 (Moon, et al., 2021). Among these, MRR was chosen as the main metric, because it can better evaluate the model’s ability to assign the positive target a ranking score that is distinguishable from other candidate targets (under the given drug and disease). MRR is calculated as follows:

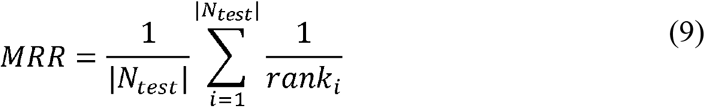

where *N*_*test*_ is the test sample set, and *rank*_*i*_ is the score ranking of the true target entity among all target candidates for the *i*_*th*_ test sample. Hits@K represents the percentage of the true target entities that appear within the top K positions of overall candidate ranking during the test phase.

### 4.3 Experiments

The main objective of our experiments is to test whether properly introducing hyperbolic space can improve the performance of Euclidean translation-based KGC methods in our task. To this end, we compared our method with the representative Euclidean translation-based method DDTE detailed in the Section 3.3. For fair comparison, we first chose FuLLiT (termed as FLONE_*base*_, later these two names will be used interchangeably), which uses self-contained drug and target embedding look-up tables, for eliminating the uncontrolled influence brought by external domain knowledge when comparing with DDTE. Then, two variants were also considered.

The first variant was named as DDTE_*bias*_, in which the drug and target type specific biases mentioned in formula (4) were added into DDTE. The second variant was the fully Euclidean counterpart of FLONE_*bas*_, termed as FEC FLONE_*bas*_ : on the top of DDTE_*bias*_, the vector-based disease offset was replaced by the matrix-based offset (i.e., using 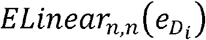 as the offset where *ELinear*_*n,n*_ and 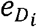 were the Euclidean linear layer and Euclidean drug embedding separately). This can eliminate the influence caused by the different calculation way (addition/multiplication) and parameter number of the disease translation offsets.

To conduct more comprehensive comparison, we also added two other types of Euclidean KGC methods ConvE (Dettmers, et al., 2018) (Convolution-based) and DistMult (bilinear product-based) (Yang, et al., 2014) as well as another hyperbolic method MuRP (Poincaré space-based) (Balazevic, et al., 2019). Furthermore, we did experiments based on two embedding dimensions: 16 and 128. 16 and 128 are representative dimensions for hyperbolic and Euclidean embedding separately (Chen, et al., 2021; Moon, et al., 2021). Theoretically, hyperbolic embedding loses less information than Euclidean embedding when embedding dimension is small. Besides, the Adam optimizer (Kingma and Ba, 2014) was adopted for all the Euclidean-based models.

The experimental results are shown in Tables 1 and 2, we can observe that, FLONE_*base*_ achieved overall better performance compared with all involved Euclidean-based models no matter on the 16/128-dimension and DTINet/BioKG, and base only under Hits@10, the sub-optimal performance was obtained on DTINet. Compared with the second-best model (based on the main metric MRR), FLONE_*base*_ obtained 5.5% and 5.8% (on DTINet) and 8.0% and 5.7% (on BioKG) performance margins on the 16-dimension and 128-dimension, respectively. The above clearly verified the effectiveness of FLONE and our hypothesis. Interestingly, MuRP did not produce very close results with our Lorentz space-based method due to its numerical instability, which indicated that Lorentz space could be more effective than the Poincaré space in our task.

**Table 1.** Comparison results of involved methods under the 16 embedding dimension.

| DTINet | MRR | Hits@1 | Hits@3 | Hits@10 |
| --- | --- | --- | --- | --- |
| $FLONE_{base}$ | **0.3806** | **0.3301** | **0.3994** | 0.4685 |
| $FEC - FLONE_{base}$ | 0.3608 | 0.3174 | 0.3680 | 0.4484 |
| DDTE<sub>bias</sub> | 0.3134 | 0.2315 | 0.3520 | 0.4651 |
| DDTE | 0.3049 | 0.2123 | 0.3518 | **0.4790** |
| MuRP | 0.3234 | 0.2463 | 0.3605 | 0.4636 |
| ConvE | 0.3160 | 0.2362 | 0.3583 | 0.4648 |
| DistMult | 0.3014 | 0.2148 | 0.3387 | 0.4739 |
| BioKG | MRR | Hits@1 | Hits@3 | Hits@10 |
| FLONE<sub>base</sub> | **0.4706** | **0.3786** | **0.5224** | **0.6452** |
| FEC – FLONE<sub>base</sub> | 0.3594 | 0.3049 | 0.3797 | 0.4600 |
| DDTE<sub>bias</sub> | 0.4222 | 0.3302 | 0.4653 | 0.5992 |
| DDTE | 0.3545 | 0.2417 | 0.4162 | 0.5586 |
| MuRP | 0.4358 | 0.3373 | 0.4911 | 0.6146 |
| ConvE | 0.3737 | 0.2815 | 0.4182 | 0.5507 |
| DistMult | 0.2877 | 0.1986 | 0.3310 | 0.4518 |

The bold data indicates the best result under current evaluation metric and dataset.

**Table 2.** Comparison results of involved methods under the 128 embedding dimension.

|  |  |  |  |  |
| --- | --- | --- | --- | --- |
| DTINet | MRR | Hits@1 | Hits@3 | Hits@10 |
| FLONE<sub>base</sub> | **0.4335** | **0.3936** | **0.4428** | 0.5054 |
| FEC – FLONE<sub>base</sub> | 0.4098 | 0.3705 | 0.4139 | 0.4907 |
| DDTE<sub>bias</sub> | 0.3893 | 0.3232 | 0.4165 | **0.5125** |
| DDTE | 0.3139 | 0.2186 | 0.3635 | 0.4997 |
| MuRP | 0.3310 | 0.2502 | 0.3699 | 0.4823 |
| ConvE | 0.3885 | 0.3389 | 0.4090 | 0.4702 |
| DistMult | 0.3555 | 0.3000 | 0.3780 | 0.4536 |
| BioKG | MRR | Hits@1 | Hits@3 | Hits@10 |
| FLONE<sub>base</sub> | **0.5027** | **0.4116** | **0.5495** | **0.6758** |
| FEC – FLONE<sub>base</sub> | 0.4549 | 0.3790 | 0.4914 | 0.5960 |
| DDTE<sub>bias</sub> | 0.4754 | 0.3780 | 0.5215 | 0.6671 |
| DDTE | 0.3806 | 0.2549 | 0.4576 | 0.6071 |
| MuRP | 0.4750 | 0.3749 | 0.5323 | 0.6586 |
| ConvE | 0.4641 | 0.3781 | 0.5090 | 0.6296 |
| DistMult | 0.3863 | 0.3176 | 0.4198 | 0.5108 |

The bold data indicates the best result under current evaluation metric and dataset.

### 4.4 Fusing domain knowledge for FLONE

After demonstrating our basic hypothesis, as described in the Section 4.2, a challenging prediction scenario where unknown drugs and targets are included was base established. It is difficult for FuLLiT/FLONE_*base*_ and DDTE to handle previously unseen drugs and targets because what they leverage is only known structural information of the DDT network. Intuitively, the fusion of (similarity) domain knowledge of involved drugs and targets could be helpful to improve the prediction accuracy of FuLLiT in this scenario. To investigate this, based on the larger and more complex DTINet DDT network and same model evaluation settings, the following variants were added into the comparison.

The first was FuLLiT with the hyperbolic drug encoder and hyperbolic target encoder using target sequence similarity (termed as FLONE_*ECFP−SEQ*_), the second was FuLLiT with the hyperbolic drug encoder and hyperbolic target encoder using GO-PPI network (termed as FLONE_*ECFP-NET*_), the third was FuLLiT with the hyperbolic drug encoder and self-contained target embedding look-up table (termed as FLONE_*ECFP-None*_), the fourth was FuLLiT with the self-contained drug embedding look-up table and hyperbolic target encoder using target sequence similarity (termed as FLONE_*None-SEQ*_), and the fifth was FuLLiT with the self-contained drug embedding look-up table and hyperbolic target encoder using GO-PPI network (termed as FLONE_*None-NET*_). The evaluation results of these variants are in Table 3.

**Table 3.** Comparison results of involved FLONE variants under the 16/128 embedding

| Methods (dim=16) | MRR | Hits@1 | Hits@3 | Hits@10 |
| --- | --- | --- | --- | --- |
| $\text{FLONE}_{\text{ECFP-SEQ}}$ | **0.4740** | **0.4121** | **0.4945** | **0.5946** |
| $\text{FLONE}_{\text{ECFP-NET}}$ | 0.4475 | 0.3841 | 0.4776 | 0.5599 |
| $\text{FLONE}_{\text{ECFP-None}}$ | 0.4322 | 0.3750 | 0.4568 | 0.5347 |
| $\text{FLONE}_{\text{None-SEQ}}$ | 0.4068 | 0.3461 | 0.4233 | 0.5260 |
| $\text{FLONE}_{\text{None-NET}}$ | 0.3865 | 0.3325 | 0.4102 | 0.4804 |
| $\text{FLONE}_{\text{base}}$ | 0.3806 | 0.3301 | 0.3994 | 0.4685 |
| Methods (dim=128) | MRR | Hits@1 | Hits@3 | Hits@10 |
| $\text{FLONE}_{\text{ECFP-SEQ}}$ | **0.5644** | **0.5139** | **0.5793** | **0.6564** |
| $\text{FLONE}_{\text{ECFP-NET}}$ | 0.5084 | 0.4606 | 0.5308 | 0.5911 |
| $\text{FLONE}_{\text{ECFP-None}}$ | 0.5033 | 0.4571 | 0.5216 | 0.5912 |
| $\text{FLONE}_{\text{None-SEQ}}$ | 0.4820 | 0.4410 | 0.4892 | 0.5614 |
| $\text{FLONE}_{\text{None-NET}}$ | 0.4363 | 0.3923 | 0.4510 | 0.5121 |
| $\text{FLONE}_{\text{base}}$ | 0.4335 | 0.3936 | 0.4428 | 0.5054 |

The bold data indicates the best result under current evaluation metric.

From the results, we found that, FLONE_*ECFP-SEQ*_ had the best performance, under 16 and 128 dimensions, it obtained 24.5% and 30.2% performance improvements on MRR base compared with FLONE_*base*_. This can demonstrate the importance of similarity domain knowledge in scenario where unknown drugs and targets occur. Interestingly, FLONE_*ECFP-SEQ*_ and FLONE_*None-SEQ*_ were better than and FLONE_*ECFP-NET*_ and FLONE_*None-NET*_ respectively. We thought the contributing reason was that, the graph-based target encoder brought unnecessary complexity, causing negative effects on the learning process, making it worse than the sequence-based target encoder. Besides, FLONE_*ECFP-None*_ was better than FLONE_*None-NET*_ and FLONE_*None-SEQ*_, which indicated that ECFP6 could be more effective compared with the provided two types of target features in our task.

### 4.5 Fusing domain knowledge for the Euclidean-based model

On the top of the conclusion from the last section, to investigate the performance gain of Euclidean translation-based models from importing domain knowledge, based on the best Euclidean model FEC - FLONE_*base*_ (based on DTINet and MRR), we did the following experiments still keeping the same model evaluation settings.

We run FEC – FLONE_*ECFP-SEQ*_, FEC – FLONE_*ECFP-None*_, and FEC - FLONE_*None-SEQ*_ on DTINet, in which all hyperbolic operations were replaced by corresponding Euclidean ones, and the meaning of subscripts of these model names was consistent with that in the last section. As shown in Table 4, FEC - FLONE_*base*_ obviously benefited from ECFP6 and target sequence similarity:

**Table 4.** Comparison results of involved Euclidean models under the 16/128 embedding dimensions.

| Methods (dim=16) | MRR | Hits@1 | Hits@3 | Hits@10 |
| --- | --- | --- | --- | --- |
| FEC – FLONE<sub>ECFP-SEQ</sub> | **0.4537** | **0.3842** | **0.4781** | **0.5958** |
| FEC – FLONE<sub>ECFP-None</sub> | 0.4312 | 0.3694 | 0.4482 | 0.5558 |
| FEC – FLONE<sub>None-SEQ</sub> | 0.4145 | 0.3630 | 0.4295 | 0.5161 |
| FEC – FLONE<sub>base</sub> | 0.3608 | 0.3174 | 0.3680 | 0.4484 |
| Methods (dim=128) | MRR | Hits@1 | Hits@3 | Hits@10 |
| FEC – FLONE<sub>ECFP-SEQ</sub> | **0.4759** | **0.4113** | **0.5006** | **0.6025** |
| FEC – FLONE<sub>ECFP-None</sub> | 0.4602 | 0.4004 | 0.4815 | 0.5775 |
| FEC – FLONE<sub>None-SEQ</sub> | 0.4519 | 0.4052 | 0.4627 | 0.5452 |
| FEC – FLONE<sub>base</sub> | 0.4098 | 0.3705 | 0.4139 | 0.4907 |

The bold data indicates the best result under current evaluation metric.

Compared with FEC - FLONE_*base*_, based on MRR, 25.7% and 16.1% performance improvements were achieved on dimensions 16 and 128, respectively. However, FLONE_*ECFP-SEQ*_ still outperformed FEC - FLONE_*ECFP-SEQ*_, which can further demonstrate the effectiveness of introducing Lorentz space embedding.

### 4.6 Extra ablation study

To confirm that the performance gain seen in FLONE having drug and target encoders compared with the one without the encoders, is not solely due to the later one’s inability to handle unseen drugs and targets, we devised an extra ablation study: We followed the same model evaluation settings, to run FuLLiT/ FLONE_*base*_ and FLONE_*ECFP-NET*_ again on DTINet. The only difference here was that we divided the original test set of each independent repeat into three parts, and computed and reported the average evaluation metrics on each part separately.

Specifically, the first part (Part 1) includes the samples corresponding to drugs that are in the original test set but not in the training set, and the second part (Part 2) includes the samples corresponding to targets that are in the original test set but not in the training set. The third part (Part 3) includes the samples corresponding to already seen drugs and targets. The numbers of unseen drugs, unseen targets, and samples for each independent repeat are in the Supplementary Material.

The results are shown in Table 5, we observed that, FLONE_*ECFP-NET*_ clearly out-performed FLONE_*base*_ on all three parts. This indicated that introducing similarity-based drug and target encoders into FLONE not only improves its predictions on triples related to seen drugs and targets, but also makes FLONE have the capability to provide effective predictions on ones corresponding to previously unseen drugs and targets.

**Table 5.** Comparison results of involved methods in the extra ablation study.

| Part Name | Methods (dim=16) | Hits@10 | Hits@3 | Hits@1 | MRR |
| --- | --- | --- | --- | --- | --- |
| Part 1 | FLONE<sub>base</sub> | 0.0596 | 0.0201 | 0.0080 | 0.0313 |
| Part 1 | FLONE<sub>ECFP-*NET*</sub> | **0.4367** | **0.3352** | **0.2379** | **0.3092** |
| Part 2 | FLONE<sub>base</sub> | 0.0000 | 0.0000 | 0.0000 | 0.0030 |
| Part 2 | FLONE<sub>ECFP-*NET*</sub> | **0.1872** | **0.1222** | **0.0779** | **0.1167** |
| Part 3 | FLONE<sub>base</sub> | 0.6473 | 0.5589 | 0.4635 | 0.5291 |
| Part 3 | FLONE<sub>ECFP-*NET*</sub> | **0.7054** | **0.5978** | **0.5111** | **0.5758** |
| Part Name | Methods (dim=128) | Hits@10 | Hits@3 | Hits@1 | MRR |
| Part 1 | FLONE<sub>base</sub> | 0.1088 | 0.0363 | 0.0140 | 0.0494 |
| Part 1 | FLONE<sub>ECFP-*NET*</sub> | **0.5587** | **0.4528** | **0.3415** | **0.4171** |
| Part 2 | FLONE<sub>base</sub> | 0.0000 | 0.0000 | 0.0000 | 0.0028 |
| Part 2 | FLONE<sub>ECFP-*NET*</sub> | **0.2016** | **0.1561** | **0.1146** | **0.1483** |
| Part 3 | FLONE<sub>base</sub> | 0.6890 | 0.6163 | 0.5522 | 0.6000 |
| Part 3 | FLONE<sub>ECFP-*NET*</sub> | **0.7650** | **0.6870** | **0.6260** | **0.6723** |

The bold data indicates the best result under current evaluation metric and tested data.

### 4.7 Visualization of the embedding spatial layout

To investigate the difference of embedding spatial layout between the Lorentz KGC model and its Euclidean counterpart, based on the 128-dimension FLONE_*ECFP-SEQ*_ and FEC - FLONE_*ECFP-SEQ*_ (on DTINet), we projected the Lorentz embeddings and Euclidean embeddings of all candidate target entities (i.e., the targets in the target entity set), under the drug entity *Nitrazepam* (*DB01595*) and disease relation *leukemia, myeloid, acute*, to the 2-dimension (Figure 3). Specifically, first the Lorentz embeddings were mapping to their isomorphic Poincaré embeddings using the formula (10), where *P*^*i*^ is the *i*_*th*_ Poincaré embedding, and 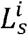 and 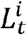 are the spatial and time dimensions of corresponding Lorentz embedding *L*^*i*^:

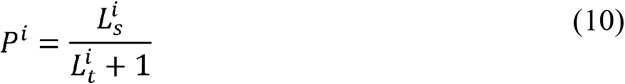

**Fig. 3.**
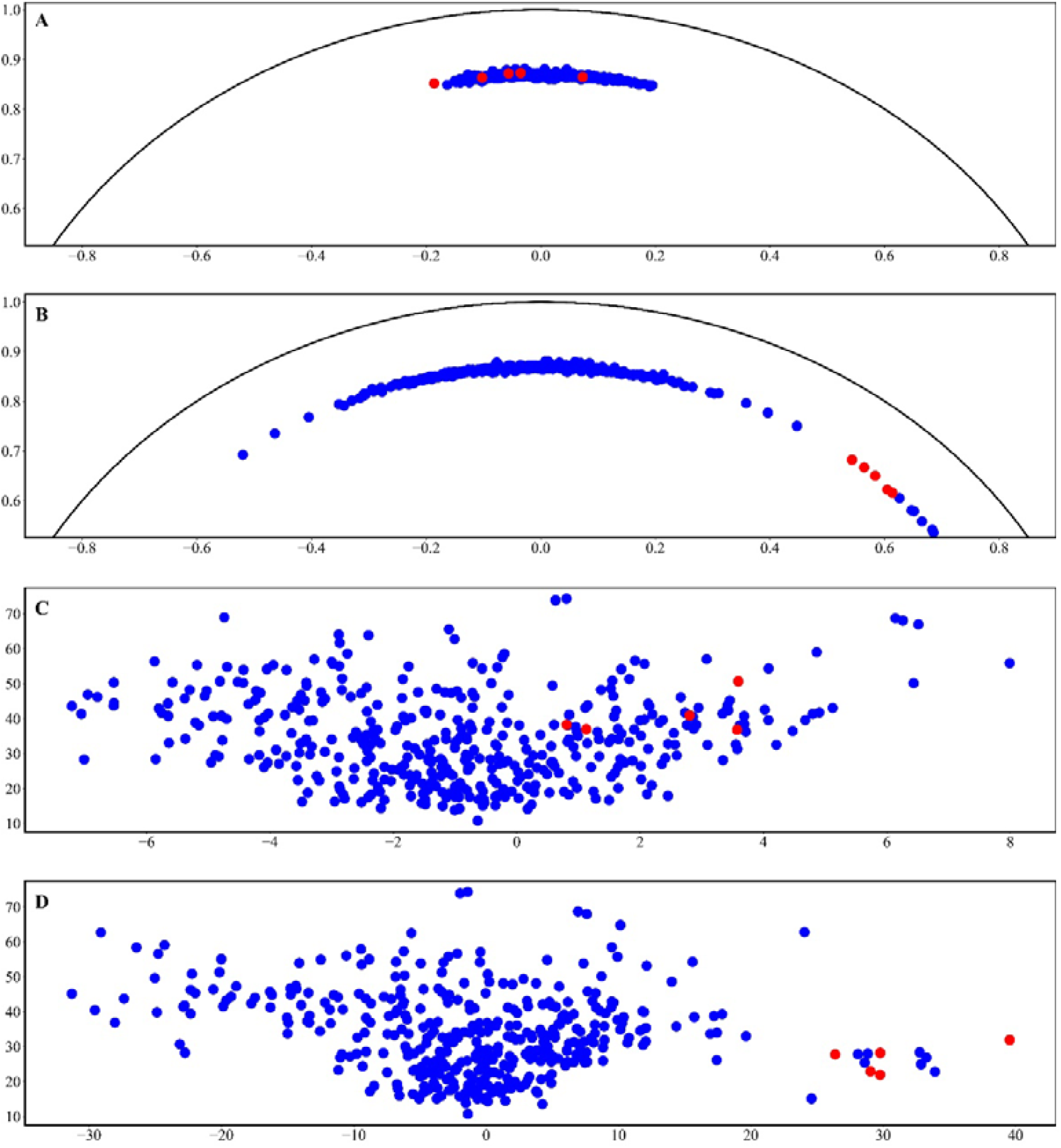
Visualization of the 2-dimension embedding spatial layout of 128-dimension FLONE_ECFP-SEQ_ and its Euclidean counterpart. These four layouts display all candidate target embeddings (under the given drug entity *Nitrazepam* and disease relation *leukemia, myeloid, acute*) in different situations (A)-(D). The blue points represent current all candidate target embeddings, and the red points represent the embeddings of current positive targets in the test set under the given drug and disease (after dimension reduction). (A) The hyperbolic target embedding layout before applying the translation of the given disease. (B) The hyperbolic target embedding layout after applying the translation of the given disease. (C) The Euclidean target embedding layout before applying the translation of the given disease. (D) The Euclidean target embedding layout after applying the translation of the given disease. Besides, the black arc in layouts (A) and (B) is the boundary of the Poincaré space.

Then, to project all high-dimensional Poincaré and Euclidean target embeddings to 2-dimension, formulas (11) - (12) by Balazevic, et al. were used, these formulas can keep the original distances and angles of these target embeddings relative to the given drug embedding (Balazevic, et al., 2019):

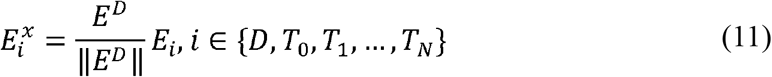

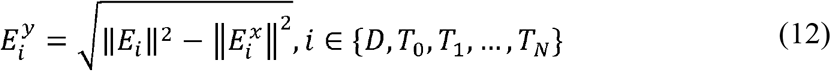

where 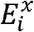 and 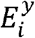 are the coordinate on the *x*-axis and *y*-axis of *i*_*th*_ new projected embedding, *E*^*D*^ is the original high-dimensional embedding of the given drug, *N* is the number of candidate target entities, and ‖ · ‖ represents the Euclidean norm.

Based on the 2-dimension spatial layout of candidate target embeddings, we found that, after applying the translation offset of *leukemia, myeloid, acute*, both FLONE_*ECFP-SEQ*_ and FEC – FLONE_*ECFP-SEQ*_ can separate the embeddings of positive targets for the given drug *Nitrazepam* (i.e., red points in Figure 3) from most of the candidate target embeddings. However, the separation of red points based on Lorentz embedding was relatively clearer. Moreover, for FLONE_*ECFP-SEQ*_, the target entities were embedded close to the boundary of the Poincaré space where the distance was larger than that around its origin, which could indicate that there is more space for embedding the entities and separating them (Balazevic, et al., 2019).

## Conclusion

This paper first hypothesizes that, because heterogeneous DDT networks could possess hierarchical structures, the translation-based KGC method can benefit from properly introducing hyperbolic space which is natural for representing network hierarchies. Within the scope of a more practical drug target prediction problem -- explicitly considering disease types and directly ranking all candidate targets, we formulated this problem as a hyperbolic translation-based triple target entity completion task to test our hypothesis. We proposed FLONE (fully Lorentz network embedding) and evaluated it on the hierarchical DDT networks after analyzing potential data leakage issues. Our experimental results showed that, FLONE generates more accurate target predictions than its Euclidean counterparts, which supports our hypothesis. We further investigated the effectiveness of Lorentz space by plotting the spatial layout of FLONE, which showed a better capacity for embedding candidate target entities. Furthermore, we found that external domain knowledge, such as drug structural and target sequence similarities can be utilized to further improve the predictive accuracy for both seen and previously unseen triples in our task.

Apart from heterogeneous DDT networks, FLONE could be applied to other complex heterogeneous networks with a hierarchical structure. However, it is worth mentioning that, apart from hierarchical structures, real-world complex networks could exhibit other types of sub-structures, e.g., the cyclic structure. In future work, we plan to introduce more non-Euclidean space, e.g., spherical space specifically for capturing cyclic structures into our framework, making it capable of adapting to different network sub-structures.

