## Supplementary Material for "FLONE: fully Lorentz network embedding for inferring novel drug targets"

**Yang Yue** is a Ph.D. student of the School of Computer Science from the University of Birmingham, UK. His research interests include bioinformatics, machine learning and data mining.

**David McDonald** received the B.Sc., M.Sc., and Ph.D. degrees from the University of Birmingham, UK, in 2015, 2016, and 2020, respectively. He now works for AIA Insights Ltd. as a Chief Technical Officer. His main research interests are drug discovery, complex network analysis, network embedding, heuristic searches and machine learning.

**Luoying Hao** is a Ph.D. student of the School of Computer Science from the University of Birmingham, UK. Her research interests include medical image analysis, video understanding, deep learning.

**Huangshu Lei** is the Chief Scientist of YaoPharma Co., Ltd. His research interests are drug discovery and organic chemistry.

**Mark S. Butler** is the Chief Technical Officer at AIA Insights Ltd. His research interests are drug discovery, natural products and antibiotics.

**Shan He** is an associate professor at the School of Computer Science, the University of Birmingham. His research interests are machine learning, evolutionary algorithms, network medicine and drug discovery.

In the Supplementary Material, we mainly introduce two sections. In the section 1, we introduce how to construct the graph-like target prior feature – the GO-PPI network, and how to process it by the fully Lorentz graph convolutional network (GCN)-based encoder for target embedding generation. In the section 2, we introduce the statistics of the unseen drugs, unseen targets, and sample number of each independent test set (mentioned in the manuscript) through tables.

1. **The GO-PPI network and its encoder**

To construct the GO-PPI network which is used to rank disease-target (gene) pairs, similar to Chen, et al [1], we first obtain gene ontology (GO) annotations of the involved targets, these GO annotations can associate involved targets with the GO terms that describe different functions of gene products. We then retrieve valid ancestral GO terms of the GO terms found by the GO annotations. A hierarchical GO network can thus be constructed based on the collected GO terms, in which the targets and GO terms are nodes, curated relationships between GO terms as well as between GO terms and targets (i.e., GO associations) are edges. This network describes the molecular function, cellular component, and biological process of the genes that encode the involved targets [2]. The protein-protein interaction (PPI) network of the involved targets is extracted from DTINet. These two networks are termed as the GO-PPI network together.

To encode this GO-PPI network for generating hyperbolic target embeddings, we propose a fully Lorentz GCN-based target encoder. Specifically, the overall Gromovs hyperbolicity $\delta$ of the GO-PPI network is 1.5, theoretically, it could also be captured by hyperbolic models. Like the Euclidean spatial GCN, the essence of Lorentz GCN is aggregating the neighboring nodes’ features/coordinates for the central node. The aggregation method used here is the Lorentzian centroid [3, 4], which is fully defined on Lorentz space and can prevent the aggregation deviating out of the Lorentz space. The aggregation is defined as follows:

$$AGG\left( x_{i}^{agg} \right):=\frac{\sum_{j\in N_{i}} a_{ij}x_{j}^{agg}}{\sqrt{-c}|\left\| \sum_{j\in N_{i}} a_{ij}x_{j}^{agg} \right\|_{\mathcal{L}}|}$$

(1)

where $N_{i}$ is the neighboring node set of node $i$ (including the self-loop), $x_{j}^{agg}$ is the input Lorentz coordinate of node $j$ in the aggregation, and $a_{ij}$ is the aggregation weight allocated to the neighbor $j$ of node $i$. For simplicity, we use the averaging aggregation weight similar to the vanilla GCN [5]. Based on this, a complete fully Lorentz GCN layer can be expressed as $AGG({FLLinear}_{m,n}\left( x_{i}^{lin} \right))$, where $x_{i}^{lin}$ is the input Lorentz coordinate of node $i$ in its enhancement of the node feature (the enhancement is based on ${FLLinear}_{m,n}\left( x \right)$).

For the GO-PPI network, GO and PPI networks are aggregated by independent fully Lorentz GCN layers, to obtain hyperbolic target embeddings under GO and PPI respectively. To fuse these two groups of target embeddings, because there are few clear definitions of the feature fusion fully operated on Lorentz space, we use our defined fully Lorentz GCN layer to aggregate every target embedding under GO with corresponding embedding under PPI, to generate the final hyperbolic target embeddings.

2. **Statistics of the independent test sets**

**Table S1.** The statistics of each independent repeat/test set in the comparison experiments.

| DTINet | Repeat 1 | Repeat 2 | Repeat 3 | Repeat 4 | Repeat 5 |
| --- | --- | --- | --- | --- | --- |
| Unseen drug number | 48 | 48 | 56 | 62 | 52 |
| Unseen target number | 41 | 53 | 59 | 53 | 51 |
| BioKG | Repeat 1 | Repeat 2 | Repeat 3 | Repeat 4 | Repeat 5 |
| Unseen drug number | 138 | 148 | 130 | 151 | 149 |
| Unseen target number | 75 | 78 | 79 | 69 | 83 |

**Table S2.** The statistics of sample numbers of the different parts in extra ablation study.

|  | Repeat 1 | Repeat 2 | Repeat 3 | Repeat 4 | Repeat 5 |
| --- | --- | --- | --- | --- | --- |
| Part 1 sample number | 3120 | 5533 | 7558 | 6580 | 5609 |
| Part 2 sample number | 3621 | 4658 | 5163 | 4911 | 4862 |
| Part 3 sample number | 26307 | 25040 | 24569 | 24477 | 24562 |
